# Optimal filtering strategies for task-specific functional PET imaging

**DOI:** 10.1101/2024.04.25.591053

**Authors:** Murray Bruce Reed, Magdalena Ponce de León, Sebastian Klug, Christian Milz, Leo Robert Silberbauer, Pia Falb, Godber Mathis Godbersen, Sharna Jamadar, Zhaolin Chen, Lukas Nics, Marcus Hacker, Rupert Lanzenberger, Andreas Hahn

## Abstract

Functional Positron Emission Tomography (fPET) has advanced as an effective tool for investigating dynamic processes in glucose metabolism and neurotransmitter action, offering potential insights into brain function, disease progression, and treatment development. Despite significant methodological advances, extracting stimulation-specific information presents additional challenges in optimizing signal processing across both spatial and temporal domains, which are essential for obtaining clinically relevant insights. This study aims to provide a systematic evaluation of state-of-the-art filtering techniques for fPET imaging.

Forty healthy participants underwent a single [^18^F]FDG PET/MR scan, engaging in the cognitive task Tetris®. Twenty thereof also underwent a second PET/MR session. Eight filtering techniques, including 3D and 4D Gaussian smoothing, highly constrained backprojection (hypr), iterative hypr (Ihypr4D), two MRI-Markov Random Field (MRI-MRF) filters (L=10 and 14 mm neighborhood) as well as static and dynamic Non-Local Means (sNLM and dNLM respectively) approaches, were applied to fPET data. Test-retest reliability (intraclass correlation coefficient), the identifiability of the task signal (temporal signal-to-noise ratio (tSNR)), spatial task-based activation (group level t-values), and sample size calculations were assessed.

Results indicate distinct performance between filtering techniques. Compared to standard 3D Gaussian smoothing, dNLM, sNLM, MRI-MRF L=10 and Ihypr4D filters exhibited superior tSNR, while only dNLM and hypr showed improved test-retest reliability. Spatial task-based activation was enhanced by both NLM filters and MRI-MRF approaches. The dNLM enabled a minimum reduction of 15.4% in required sample size.

The study systematically evaluated filtering techniques in fPET data processing, highlighting their strengths and limitations. The dNLM filter emerges as a promising choice, with improved performance across all metrics. However, filter selection should align with specific study objectives, considering factors like processing time and resource constraints.

## Introduction

Functional Positron Emission Tomography (fPET) has become a powerful tool, enabling researchers and clinicians to investigate the intricate details of biological processes at the molecular level *in vivo*^1, 2^. This capability to visualize and quantify task-driven dynamic changes in molecular activity has far-reaching implications for understanding brain function, disease progression, and monitoring treatment efficacy. Although research into fPET began almost a decade ago^3, 4^, rapid methodological advances in the field have led to improvements in temporal resolution. Commencing with more conventional PET framing times of 60 s, fPET quickly progressed to 30 s^1, 2, 5^ and more recently 16 s^6^. Notably, a recent study unveiled acute temporal changes in the fPET signal after task performance using 3 s frames, which was previously indiscernible at lower temporal resolutions^7^. This breakthrough was achieved, in part, through specialized filtering techniques designed to enhance the temporal signal to noise ratio (tSNR), while preserving the acute task-specific changes. Given the inherently low tSNR of PET and the recent interest and rapid advances in high temporal resolution imaging, the demand for enhancing both spatial and temporal resolution in fPET data is becoming increasingly imperative. However, the potential of high-resolution fPET is fundamentally limited by challenges related to spatial^8^ and temporal resolution^9^. The precise localization of molecular events and the ability to capture rapid changes over time are essential for extracting clinically accurate and meaningful information.

Despite the considerable progress in the methodology behind fPET, achieving optimal spatial and/or temporal SNR (tSNR) remains an ongoing challenge. The need for improved resolution is emphasized by the complexities of biological systems, where rapid and subtle changes are often the key indicators of physiological and pathological processes. As such, the investigation and optimization of filtering techniques used to enhance spatial and temporal resolution in fPET is crucial for unlocking its full potential.

Existing methods to enhance fPET data predominantly involve 3D filtering techniques. The most common and widely used approach is Gaussian smoothing, employed not only in fPET^10,11^, but also in structural and functional magnetic resonance imaging (fMRI)^12, 13^. Other 3D filtering techniques exist, such as highly constrained backprojection (hypr)^14^, static non-local means^15^ (sNLM) and a more recent technique incorporating anatomical knowledge to enhance the image using a Bowsher-like prior^16^. Each of these techniques has its own advantages and limitations. Although, NLM filtering including the temporal domain has been explored in fMRI^17^ and in PET for denoising low-count frames^18^ its rigorous evaluation in the context of fPET is missing. Other techniques like Gaussian smoothing and hypr have seen recent improvements by incorporating a temporal component^19–21^.

While each filtering technique has been shown to improve spatial and/or tSNR when compared to a standard approach using either simulations or phantom data, a comprehensive assessment of the most well-known techniques on the same real-world dataset using multiple test metrics, has yet to be performed. The primary objective of this study was to assess the efficacy of various filtering techniques in optimizing fPET task imaging results, with a specific focus on practically relevant parameters such as test-retest reliability, temporal SNR, group level spatial effects and sample size estimation. By systematically evaluating and comparing different filtering techniques in a practical manner, we aim to identify the most promising approaches for enhancing the quality of fPET images.

## Materials and Methods

For this retrospective analysis, data from our previous test-retest study was used ^22^. Thus, full details about the study design and data acquisition can also found there and in related work^1,11^.

### Participants

Data from 40 healthy participants (20 male, mean age ± SD: 23.0 ± 3.4 years, all right-handed) were used, and for 20 participants, data for test-retest reliability analysis was available (10 male, 23.1 ± 3.1 years). All participants underwent a routine medical investigation at the screening visit including electrocardiography, blood tests, neurological and physiological examination, and a urine drug test. Psychiatric disorders were ruled out with the Structural Clinical Interview DSM-IV conducted by an experienced psychiatrist. Female participants additionally underwent a pregnancy test at the screening visit and before each PET/MRI measurement. Participants had to fast for at least four hours prior to the scan, including the consumption of sweetened beverages and caffeine^23^. Exclusion criteria included current or previous neurological, somatic or psychiatric diseases, current breastfeeding or pregnancy, left-handedness, substance abuse, MRI contraindications, past participation in a study with ionizing radiation exposure and regular experience playing puzzle games i.e. Tetris® or similar. Due to radiation protection, participants above 100 kg were also excluded. After a detailed explanation of the study protocol, all participants gave written informed consent. Participants were insured and reimbursed for their participation. The study was approved by the Ethics Committee of the Medical University of Vienna (ethics number: 1479/2015) and all procedures were carried out in accordance with the Declaration of Helsinki. The study was registered at ClinicalTrials.gov (ID: NCT03485066).

### Cognitive task

Participants were required to play an adapted version of Tetris®, including 2 levels of difficulty to induce various levels of cognitive load. Participants were required to perform all actions using only their right hand. To familiarize themselves with the task and the controls, participants completed a 30 s training of each condition before each scanning session. A more detailed description of the task can be found in^22, 24^.

### Study design

The experimental protocol aimed to assess cognitive tasks using a combination of continuous task performance and a conventional block design. The session started with the acquisition of a structural T1-weighted image. Thereafter, [^18^F]FDG was administered in a bolus + constant infusion protocol. The initial baseline was 8 minutes of rest. Participants then performed four task conditions of varying difficulty (6 min each, 2 easy, 2 hard in a pseudo-randomized order), followed by a 5-minute rest condition where subjects were instructed to look at a crosshair and let their thoughts wander. Additional data acquired for different purposes were not used in this study (ASL, BOLD and DTI sequences). The total scan time was 100 minutes, representing a typical duration for PET studies. See Hahn et al. for a more detailed description^1^. Data for test-retest analysis were acquired 4.2 ± 0.7 weeks after the first scan.

### Data acquisition and blood sampling

The radiotracer [^18^F]FDG was synthesized each measurement day at the Department of Biomedical Imaging and Image guided Therapy, Division of Nuclear Medicine, Medical University of Vienna. Simultaneous to fPET start, [^18^F]FDG was administered via a cubital vein as a 1 minute bolus followed by constant infusion for 51 minutes with an infusion pump (Syramed mSP6000, Arcomed, Switzerland, dosage: 5.1 MBq/kg, bolus speed: 816 ml/h, infusion speed: 42.8 ml/h, bolus-infusion ratio of activity: 20:80%), which was placed in an MR-shield. fPET data was acquired in list-mode using a Siemens 3T mMR scanner (Erlangen, Germany), enabling the retrospective definition of frame lengths during reconstruction.

A T1-weighted structural image was acquired with a magnetization prepared rapid gradient echo (MPRAGE) sequence prior to radiotracer administration (TE/TR=4.21/2200 ms, voxel size=1×1x 1.1 mm, matrix size=240×256, slices=160, flip angle=9°, TI=900 ms, 7.72 min). The image was used to rule out severe structural abnormalities, for attenuation correction^25^ and spatial normalization to MNI space.

Prior to each PET/MRI measurement, the individual fasting blood glucose level was measured as an average of a triplicate. Arterial blood samples were drawn from a radial artery throughout the radiotracer administration (time points: 3, 4, 5, 14, 25, 36 and 47 min after infusion start) and were timed not to interfere with task performance and the MRI acquisition. Blood samples were processed as previously described ^11^. In short, whole blood activity and plasma activity after centrifugation were measured in a gamma-counter (Wizard2, 3”; Perkin Elmer, USA). The whole blood curve was linearly interpolated and resampled to match the time points of the reconstructed fPET frames. The plasma to whole-blood ratio was averaged across time points. The whole blood curve was then multiplied with the mean plasma-to-whole-blood ratio to obtain an arterial input function for absolute quantification.

### Preprocessing and quantification

All fPET data was reconstructed using an Ordinary Poisson – Ordered Subset Expectation Maximization Algorithm (OP-OSEM), set at 3 iterations and 21 subsets. The output image contained a matrix size of 344 x 344 with 127 slices and a voxel size of 2.09 x 2.09 x 2.03 mm. The reconstructed data was binned into 104 frames of 30s each. Standard corrections, including dead time, decay, and scatter, were applied, and attenuation correction was performed using a pseudo-CT approach based on the structural MRI acquired during the initial measurement^25^.

Preprocessing and quantification followed established procedures from previous studies^1, 4, 5^. Specifically, SPM12 (https://www.fil.ion.ucl.ac.uk/spm) was utilized for head movement correction (quality = best, registered to mean image), spatial normalization to MNI space using the structural MRI. The mean PET image was coregistered to the structural MRI, and the resulting transformations were applied to the dynamic fPET data. Once normalized, the data was smoothed using different techniques, see filter assessment below. The smoothed data was subsequently masked using a grey matter mask (SPM12 tissue prior, thresholded at 0.1), and a low pass filter with a cutoff frequency of half the task duration was applied to the time course of each voxel.

A general linear model (GLM) was employed to distinguish between task-specific and baseline metabolism, using four regressors: baseline, easy task block, hard task block and the first principal component of the six movement regressors obtained during movement correction^4^.

The Gjedde-Patlak plot was employed to derive the influx constant (K_i_), with linearity assumed after 15 min after tracer application. This resulted in three separate K_i_ maps for rest, easy, and hard. Finally, the cerebral metabolic rate of glucose (CMRGlu) was quantified using the lumped constant of 0.89^26^. For a more detailed description please see^22^.

### Filter assessment

In this work we aimed to assess the most recognized PET filtering techniques and were implemented in MATLAB unless otherwise specified. These filters include:

#### 3D Gaussian filter

Is the most commonly used spatial smoothing technique in image processing. It operates by convolving the image with a three-dimensional Gaussian kernel. The convolution process assigns a weighted average to each voxel in the image, with the weights determined by the Gaussian distribution. This smoothing helps reduce noise and emphasize larger-scale features in the data^27^. Here we used the SPM function with a smoothing kernel with a full width half maximum (FWHM) of 8 mm.

#### 4D Gaussian filter

which extends on the concept of the 3D Gaussian filter to four dimensions, incorporating time as an additional dimension. This filter is particularly relevant in the context of dynamic imaging data, such as functional imaging over time. A spatial kernel of 8 mm FWHM and a temporal window of 5 frames (2 before and 2 after selected frame; 2.5 min) were used.

#### Hypr filter

is an advanced filtering technique designed to improve spatial resolution in medical imaging. It operates by incorporating constraints into the back-projection process during image reconstruction or preprocessing. These constraints, derived from both the acquired data and prior knowledge, guide the reconstruction algorithm to produce images with enhanced spatial details. Hypr is particularly useful in scenarios where high spatial resolution is crucial, such as in the case of small anatomical structures^14^. The composite (i.e., temporally summed) image was smoothed using an 8 mm FWHM kernel.

#### Iterative 4D hypr (Ihypr4D) filter

represents an extension of the hypr filter combined with a 4D Gaussian filter, which incorporates both spatial and temporal constraints, while the iterative step reduces errors during the filtering process through multiple iterations^21^. Parameters were selected from^19^. In summary, the segmentation of homogenous regions from the mean fPET image was performed using k-means clustering where k was set to 30 for feature extraction prior to running ihyper4D. The number of iterations were set to 4 and the smoothing kernel was set using the same parameters as the 4D Gaussian smoothing.

#### MRI-Markov Random Field (MRI-MRF)

The MRI-MRF prior, a modified Bowsher-like technique, enhances fPET image quality by incorporating anatomical information from coregistered MRI. This technique utilizes a continuous weighting scheme, patch-based similarity, and smoothly-decaying function which contribute to improved identification of brain activations^16^. Here we tested two different MRF neighborhood width parameters L = 10 mm and 14 mm, as used in^16^. Where the neighborhood refers to a spatial arrangement of nearby voxels around a central voxel.

#### sNLM Filter

is an extension of the traditional non-local means filter, adapted for time-series imaging data. The non-local means approach involves averaging pixel values based on similarity patterns in the image. In the sNLM, this concept is extended to capture temporal correlations over the entire time course in addition to spatial similarities. By considering both spatial and temporal information, the sNLM filter aims to preserve fine details in the data while effectively reducing noise in dynamic imaging sequences^15^. The following parameters were used: a search window of D = 11 voxels and a patch size of 3 x 3 x 3 voxels^18, 28^. A post smoothing 5 mm FWHM was used for better comparison to the other techniques, as this yields a total filter kernel of approximately 8 mm FWHM.

#### dNLM filter

is a further extension of the sNLM by capturing only correlations over a short duration, similar to a sliding window approach. The dNLM filter aims to preserve acute (e.g., task-induced) signal changes while reducing noise^7^. To incorporate a temporal parameter we used the standard setting as for the sNLM filter but added a 4^th^ dimension to the patch size of 5 frames or 2.5 min. Here a 5 mm FWHM post smoothing was also used.

### Statistical analysis

A rigorous analysis of the acquired data was conducted employing a multifaceted approach to ensure comprehensive insights of each filter’s properties of both spatial and temporal dimensions. For all tests three regions of interest (ROIs) were selected from previous analyses on the same dataset, which represent a robust task-activation assessed via a conjunction of three imaging modalities^1^. These regions comprise the frontal eye field (FEF), intraparietal sulcus (IPS) and the occipital cortex (OCC).

The Intraclass Correlation Coefficient (ICC) was utilized to assess the reliability and consistency of the observed effects between two repeated fPET measurements. Additionally, the individual tSNR was computed to gauge the robustness of the signal over time. This provides information about the ability to identify stimulation-induced changes in the presence of noise and, thus, an insight into the temporal information captured by the filtering techniques. At the group level, peak and mean task-specific t-values were compared to quantify the strength of neural responses and model fit after each filter technique. T-tests were corrected for multiple testing using Gaussian random field theory as implemented in SPM12 and the threshold for significance was set at p < 0.05 family-wise error (FWE)-corrected at the cluster-level following p < 0.001 uncorrected at the voxel-level and separately at the peak FWE-corrected p < 0.05. A power analysis was conducted to determine the requisite sample size for detecting a significant effect in a prospectively planned study (one sample case, α = 0.05, Power = 0.95, two-tailed). The effect size for each filtering approach was assessed by utilizing the mean and standard deviation of task-induced CMRGlu clusters, while accounting for multiple comparisons, i.e., number of voxels, via the Bonferroni adjustment method (p < 2.585* 10^-7^). CMRGlu clusters were determined using each filter’s quantified fPET maps where an overlap with the aforementioned three specified ROIs occurred. The 3D Gaussian filter was used as a reference for filter comparisons, since it represents the most commonly employed approach in functional neuroimaging.

## Results

An overview of each filter’s performance for each test can be found in table 1. A more detailed overview of each filter’s regional performance can be found in table S1-4. Group activation maps at cluster level correction (Figure 1) and voxel level correction (Figure 2) for each filter were created. While both Gaussian filtering techniques exhibit very low runtime on our data (mean runtime: ∼30 s), the MRI-MRF (mean runtime: 14 – 16.5 h) and NLM (mean runtime: 7.9 h) filters require substantially longer processing times. The NLM filters were processed on a single core for fair comparison, although parallelization is available, halving the runtime per core. The hypr and Ihypr4D filter displayed moderate mean runtimes of 4 and 5 min per dataset, respectively.

**Table 1:**
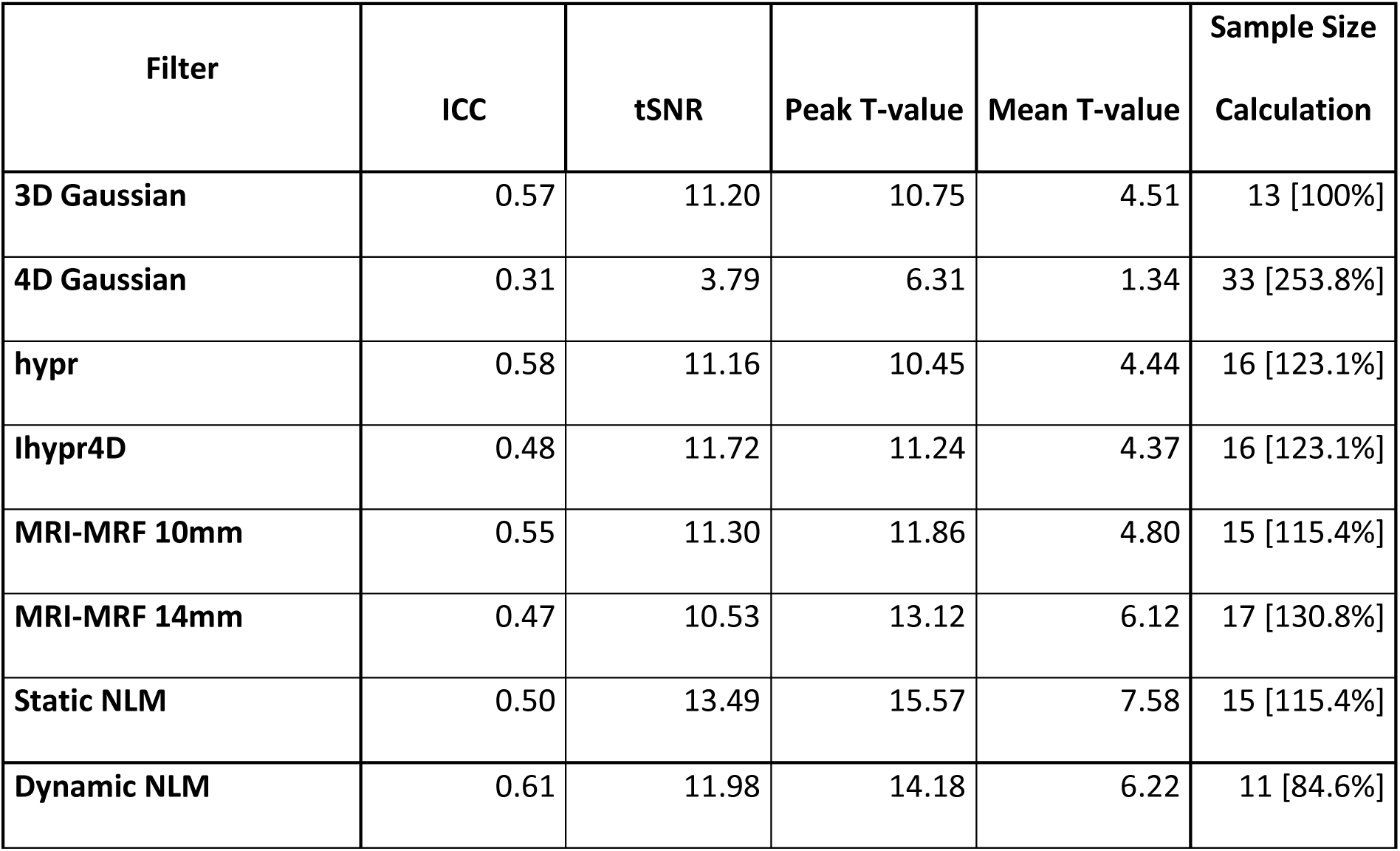
Summarized overview of all examined parameters including Intraclass Correlation Coefficients (ICC), temporal signal to noise ratio (tSNR), peak and mean T-values for each of the filtering techniques. Furthermore, the estimated sample size per filter is also listed as absolute numbers and relative to the 3D Gaussian filter in %. The values represent an average of both difficulty levels and all three regions of interest encompassing the frontal eye field, intraparietal sulcus and the occipital cortex. Separate values for each brain region and task difficulty are available in supplementary tables 1-4.

| Filter | ICC | tSNR | Peak T-value | Mean T-value | Sample Size<br>Calculation |
| --- | --- | --- | --- | --- | --- |
| **3D Gaussian** | 0.57 | 11.20 | 10.75 | 4.51 | 13 [100%] |
| **4D Gaussian** | 0.31 | 3.79 | 6.31 | 1.34 | 33 [253.8%] |
| **hypr** | 0.58 | 11.16 | 10.45 | 4.44 | 16 [123.1%] |
| **lhypr4D** | 0.48 | 11.72 | 11.24 | 4.37 | 16 [123.1%] |
| **MRI-MRF 10mm** | 0.55 | 11.30 | 11.86 | 4.80 | 15 [115.4%] |
| **MRI-MRF 14mm** | 0.47 | 10.53 | 13.12 | 6.12 | 17 [130.8%] |
| **Static NLM** | 0.50 | 13.49 | 15.57 | 7.58 | 15 [115.4%] |
| **Dynamic NLM** | 0.61 | 11.98 | 14.18 | 6.22 | 11 [84.6%] |

**Figure 1:**
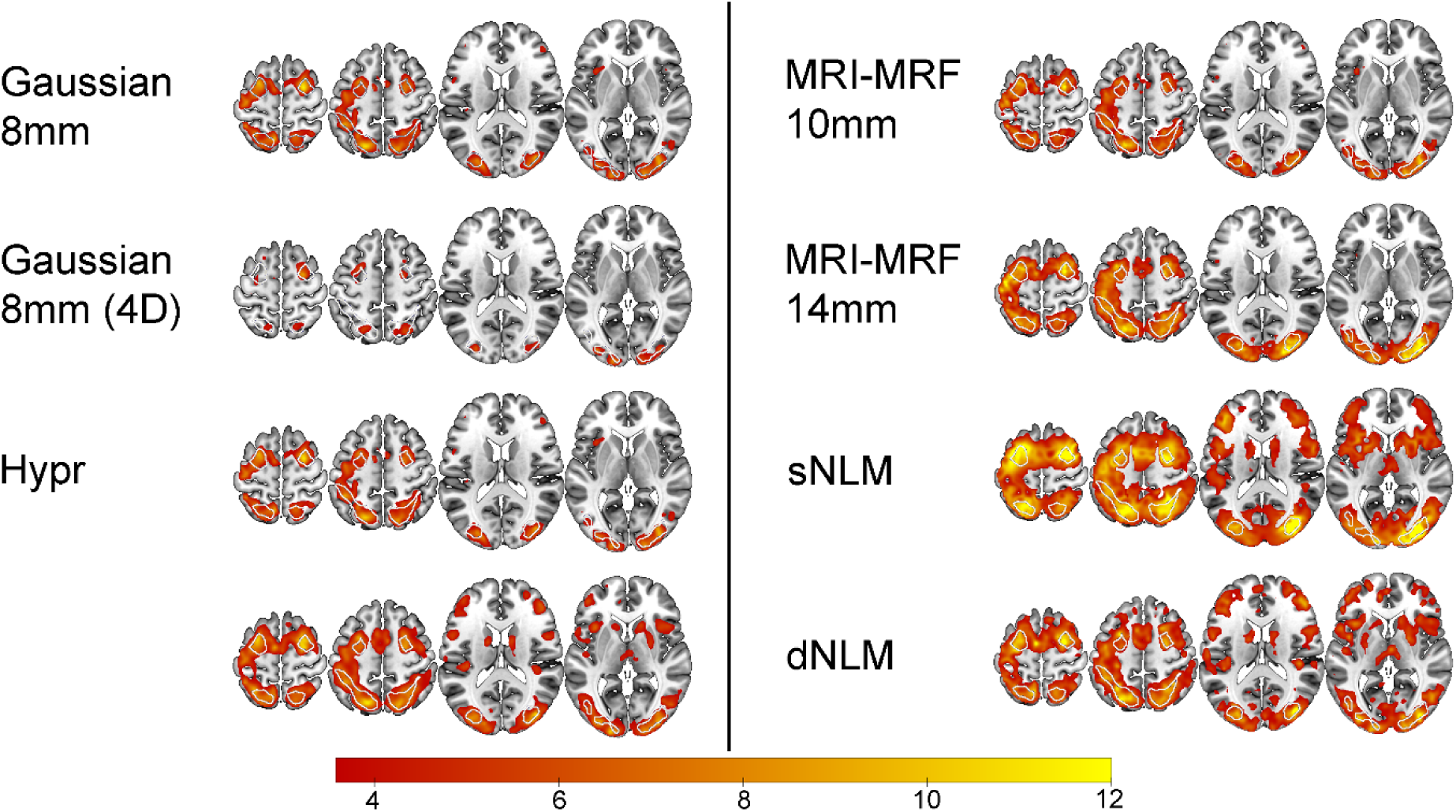
Spatial overview of task-performance (Hard > baseline) after filtering was performed. Group level statistical analysis was carried out for the hard task condition (n=40), corrected at p < 0.05 FWE cluster level, after p < 0.001 uncorrected voxel level. White outlines indicate the 3 task-active regions. The color bar represents the mean T-values.

**Figure 2:**
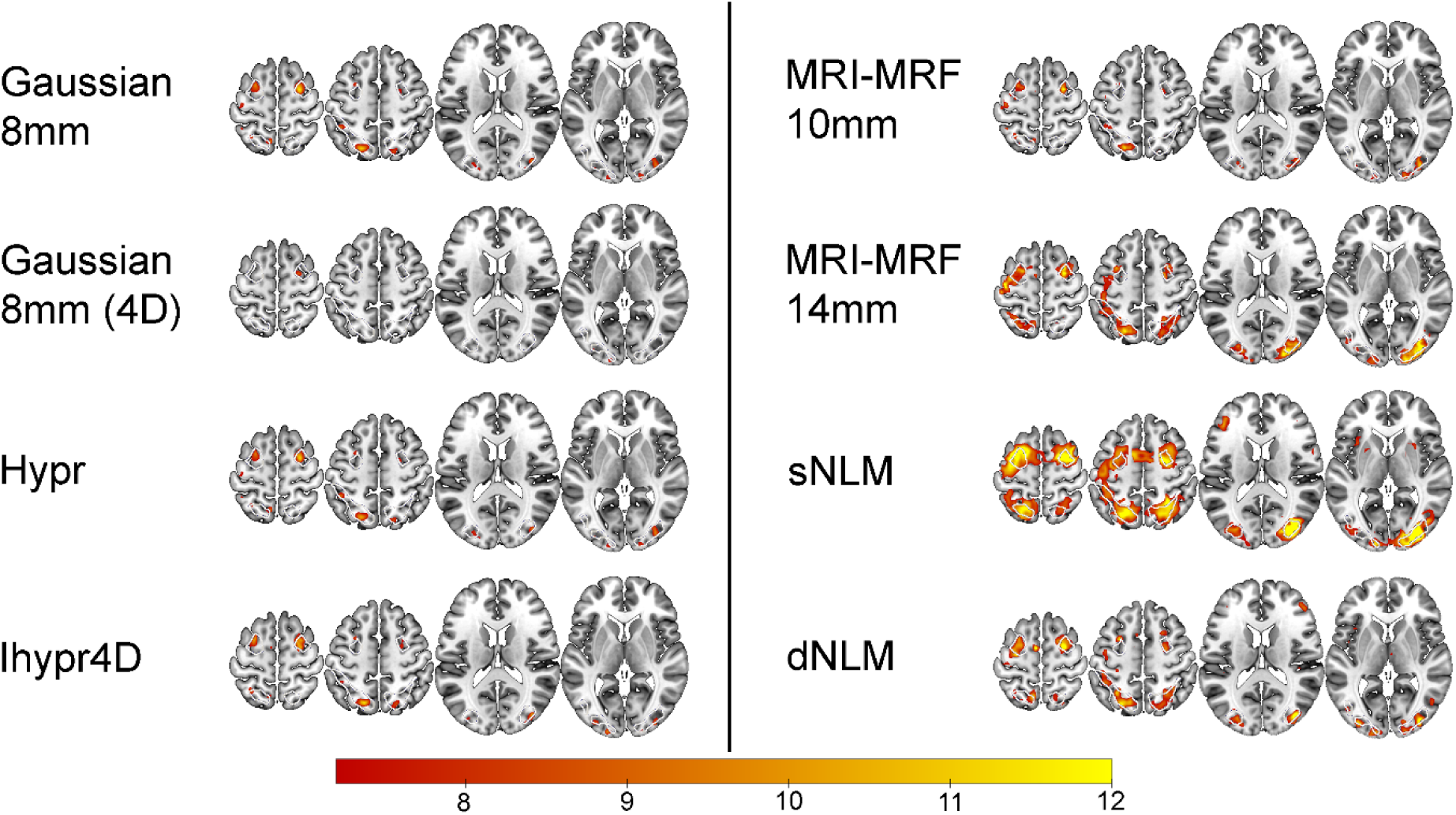
Spatial overview of task-performance (Hard > baseline) after filtering was performed. Group level statistical analysis was carried out for the hard task condition (n=40), corrected at p < 0.05 FWE voxel level. White outlines indicate the 3 task-active regions. The color bar represents the mean T-values.

### Test-retest reliability

When comparing ICC values of the 3D Gaussian filter to those obtained with other filters, only the dNLM and hypr filter showed improvements to task-based test-retest reliability. The 4D Gaussian filter, MRI-MRF L = 14 and Ihypr4D displayed the greatest decrease in ICC compared to the 3D Gaussian filter (table 1, S1). The other filter techniques (MRI-MRF L=10, sNLM) performed similarly well.

### Temporal signal-to-noise ratio

All filter approaches, excluding the 4D Gaussian smoothing, and MRI-MRF L = 14, displayed similar or improved tSNR values when compared to the 3D Gaussian smoothing (table 1, S2), whereas, the greatest improvements were seen in both NLM filtering techniques.

### Spatial task-based activation

Similar to the individual tSNR, most filter techniques displayed an improvement in group-level peak and mean t-values extracted from the task-active ROIs. The 4D Gaussian showed substantially decreased performance compared to the 3D Gaussian filter, whereas, the hypr indicated similar t-values. Increased mean and peak t-values were found for both MRI-MRF and highest values were observed for both NLM filtering techniques (table 1, S3-4). Task-based spatial activation maps for each filtering technique can be found in figure 1 and 2.

### Temporal differences

The temporal dynamics of each filtering technique were scrutinized through the analysis of the task-specific fPET signal, as illustrated in figure 3, supplementary figures 1 and 2. Notably, the MRI-MRF 14 and Gaussian 4D filters exhibited a tendency to overly smooth the signal, which is in line with the lower tSNR for these two filters (table 1). The MRI-MRF 14, while capturing more acute changes in the PET signal, tended to excessively smooth out the overall task trend (Figure 3). Conversely, the Gaussian 4D filter demonstrated an opposite effect, preserving the general task effect but smoothing the acute changes excessively (figure 3). In contrast, the hypr, and MRI-MRF 10 filters demonstrated a better temporal alignment with the Gaussian 3D signal, showing reduced peaks while maintaining signal stability, see figure 3. The sNLM filter exhibited a notably smoother time course compared to the hypr, and MRI-MRF 10 filters, yet still followed the general time course of the Gaussian 3D data. The dNLM filter, akin to the sNLM, presented a smoother time course while better preserving the amplitude of task-induced changes. Interestingly, the MRI-MRF 14 was observed to underestimate tracer uptake also for the baseline condition when compared to other filtering techniques, as depicted in supplementary figure 1.

**Figure 3:**
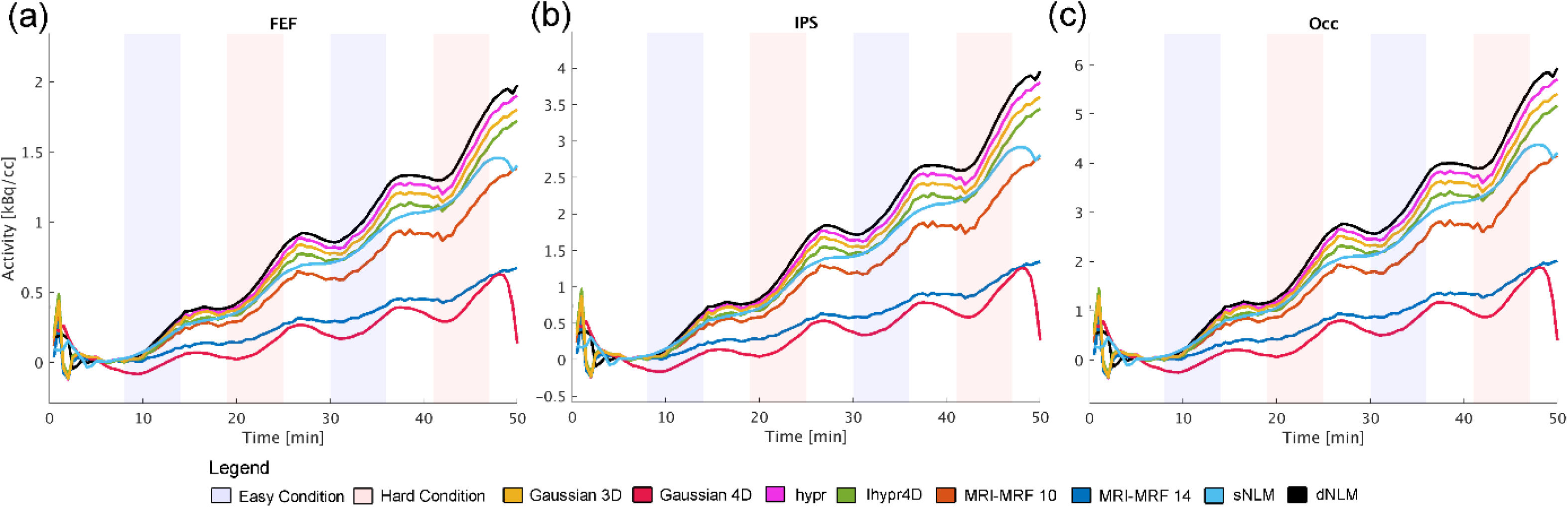
Illustration of the average task regressors’ time course for each filtering technique for a subset of participants whose task difficulty was ordered as easy-hard-easy-hard (n = 10), extracted from each task-positive region of interest. Excluding the MRI-MRF 14 filter, each filter successfully captured the task-induced increases in glucose metabolism. The MRI-MRF 14 (dark blue) and 4D Gaussian (red) filters exhibited a notable detrimental impact on baseline uptake, evident in the flatter lines, pronounced to a much lesser degree in the MRI-MRF 10 (orange). Filters such as hypr (pink), Ihypr4D (green), and dNLM (black) displayed uptake profiles most akin to the 3D Gaussian filter (yellow), with dNLM demonstrating the least noise. Similarly, sNLM (light blue) exhibited a high signal-to-noise ratio akin to dNLM, albeit with attenuated task-specific effects. These regions included the (a) frontal eye field (FEF), (b) intraparietal sulcus (IPS) and (c) the occipital cortex (OCC).

### Sample size calculation

The power analysis revealed varying sample size requirements across filter techniques, with the 3D Gaussian filter necessitating 13 participants to detect a meaningful effect. In comparison, hypr and Ihypr4D both required 16 participants, representing 123.1% of the sample size needed for 3D Gaussian. Similarly, MRI-MRF 10, MRI-MRF 14 and sNLM required 15, 17, and 15 participants, respectively, reflecting 115.4%, 130.8%, and 115.4% of the reference sample size. The 4D Gaussian required 33 participants (253.8%), while the dNLM exhibited the lowest requirement with 11 participants or 84.6% of the 3D Gaussian sample size, see table 1 for an overview.

## Discussion

The aim of this paper was to systematically evaluate various filtering techniques in the context of stimulation-induced changes in glucose metabolism using fPET, with a focus on practically relevant parameters such as test-retest variability, tSNR, and spatial task-based activation. The results present a nuanced understanding of the strengths and weaknesses of each filtering method, providing valuable information to choose the most suitable technique.

### Spatial filtering techniques

The comparison of 3D Gaussian, hypr, sNLM and MRI-MRF 10 filters revealed similar performance across all metrics. These similarities, coupled with the additional complexity and processing time of the sNLM and MRI-MRF filter, emphasize the importance of considering performance and computational efficiency. While the MRI-MRF filter technique shows decent task-induced activation at the group level, it faces challenges due to the absence of a temporal component^16^. The crucial parameter L (i.e., MRF neighborhood) in MRI-MRF was found to significantly influence its performance, introducing a trade-off between spatial resolution and overall efficacy. When choosing the L parameter, the decision should be guided by the specific requirements of the study, as it has a profound impact on the spatial resolution. While the sNLM showed high spatial task-based activation statistics, its test-retest variability remained average when compared to other spatial filtering techniques. Upon examining the spatial task-based activation (figure 1 and 2), it appears that the elevated t-statistic values (table 1) may result from the overestimation of task-based activation (see below, section: Temporal and spatial differences). This further shows that increasing effect sizes do not always signify favorable outcomes when employing certain filtering techniques, as they may also indicate overfitting or other undesirable effects.

### Spatiotemporal filtering techniques

While the 4D Gaussian, Ihypr4D and dNLM filter techniques employ a similar sliding-window approach with comparable spatial and temporal parameters, pronounced differences in the outcome parameters were found. The 4D Gaussian filter exhibited suboptimal performance in all categories, as evidenced by its decreased uptake during task and rest, which could be attributed to excessive temporal smoothing over the task effect. This highlights the importance of preserving temporal features. In contrast, the dNLM filter, which emerged as a more robust choice, employs a local search window and patch-based strategy^18^. This allows for the preservation of task-induced features while adapting to variations in temporal signals. Application of such an adaptive smoothing kernel, as seen in NLM, is further supported by its ability to maintain sharp edges, by not sampling voxels outside the brain or grey matter. The Ihypr4D filter showcased improved results when compared to the 4D Gaussian filter, which could be attributed to its iterative processing and inclusion of a composite image.

### Implications of filter selection

While every study poses a unique hypothesis, it is important to note that each filter has its strengths and weaknesses. The output parameters tested highlight their relevance in different facets of fPET analyses. The emphasis on tSNR, which is important for single subject analysis and comparing task conditions, highlights the superior performance of the Ihypr4D, MRI-MRF L=14, sNLM, and dNLM filters. On the other hand, power calculations, which are usually performed using task-based activations (i.e., peak or mean T-values), favored static and dynamic NLM filters, while MRI-MRF also demonstrated decent improvements compared to standard 3D Gaussian filtering. The assessment of ICC for comparing the reliability between multiple measurements emphasized the strengths of hypr and dNLM filters. Albeit with the caveat of the hypr filter not taking the temporal component into account, which can be crucial when working with high-temporal resolution fPET data^7^. Furthermore, the observed variations in sample size requirements across different filtering techniques underscore the impact of methodological choices on statistical power. The 4D Gaussian demonstrated notably lower power, necessitating larger sample sizes compared to the 3D Gaussian filter, which implies increased resource allocation. On the other hand, the hypr, Ihypr4D, MRI-MRF 10 and 14 as well as sNLM exhibited comparable sample size requirements. Notably, the dNLM stood out with the lowest sample size demand, potentially decreasing the sample size needed for a study. Consequently, this reduction may play a vital role when disentangling more nuanced task effects between different conditions. The selection of an appropriate filter, therefore, becomes contingent on the specific metric of interest, underlining the need for a nuanced approach based on the study’s objectives.

### Temporal and spatial differences

The temporal overview of various filtering techniques offers critical insights into their effects on the fPET signal dynamics. The observed tendency of the MRI-MRF 14 and Gaussian 4D filters to excessively smooth the signal in the temporal domain highlights potential drawbacks in capturing both acute changes and the general task effect. Conversely, the Gaussian 3D, hypr, and MRI-MRF 10 filters demonstrated a more balanced approach, reducing peaks while maintaining signal stability. The sNLM filter demonstrated a notably smoother global time course, suggesting a potential noise reduction without compromising the overall signal pattern observed in the data. However, this excessive temporal smoothing, driven by the large temporal window, appears to be influenced more by baseline uptake, which is considerably higher, rather than the task-induced uptake changes. Additionally, these factors contribute to an overestimation of task activation throughout the brain and have been demonstrated to dampen the magnitude of task-induced effects. Consequently, this could result in a lack of spatial specificity, despite producing high task-specific group statistic metrics. The dNLM filter further underscored the ability to maintain the amplitude of task-induced changes, while providing a time course with less noise. Moreover, the MRI-MRF 14’s underestimation of both global and task-active tracer uptake indicates that using a large neighborhood parameter (L = 14) is not optimal for fPET. The MRI-MRF 10 shows a more similar tracer uptake when compared to other filtering techniques. Furthermore, both NLM techniques and MRI-MRF 14 show markedly improved T-values at peak level and across the entire task-active region in comparison to other filter techniques. All in all, filtering techniques that overly smooth the time course are not suitable in high temporal resolution fPET frameworks^7^.

### Processing time

A consideration of processing time provided valuable insights into the feasibility of implementing different filters on typical hardware configurations. While 3D Gaussian smoothing proved to be a quick and efficient option, the high processing times associated with NLM and MRI-MRF filters or the requirement for dedicated processing servers underlined the need for careful consideration in resource-constrained environments. The hypr, Ihypr4D, and 4D Gaussian smoothing fell within a moderate processing time range, making them more practical for a broader range of applications and hardware options. Although all filters were programmed in MATLAB, optimizing the code or utilizing other programming languages capable of more efficient data processing could significantly decrease the runtime of complex filtering techniques, potentially rendering them more suitable for general use.

### Limitations

While the absence of a ground truth method can be seen as a limitation, we relied on the 3D Gaussian filter as a reference, due to its widespread usage in multiple modalities. Furthermore, the definition of task-active ROIs using a conjunction of signals may introduce a potential source of bias, but on the other hand makes it less dependent on fPET alone. The limited sample size, although typical for PET studies, poses constraints, for the future assessment of advanced techniques such as deep learning^29, 30^, where larger datasets are often required. The use of standard algorithm parameters from previous literature, while practical, may not capture the full spectrum of parameter space, limiting the generalizability of the findings. However, an exhaustive evaluation is limited by the computational expense of several filters. The structural component was not included in filters other than MRI-MRF, representing another limitation. This could potentially influence the overall performance of the filters, particularly in scenarios where structural information is crucial.

## Conclusion

We aimed to provide an overview of the strengths and limitations of various filtering approaches in the context of fPET studies. The choice of filtering technique should be tailored to the specific parameter of interest in improving the hypothesis. The dNLM filter emerges as a promising compromise, exhibiting the best overall performance across various metrics. However, it may be less suitable for specific use cases such as when minimal processing power is available or only a specific parameter is of interest. Following closely are the MRI-MRF L = 10 and the hypr filters, each offering unique advantages. The 3D Gaussian filter stands out for its efficient processing time and respectable performance, still making it a viable option in scenarios where computational efficiency is paramount.

## Acknowledgments

This research was funded in whole or in part by the Austrian Science Fund (FWF) [grant DOI: 10.55776/KLI610, PI: A. Hahn]. For open access purposes, the author has applied a CC BY public copyright license to any author accepted manuscript version arising from this submission. S. Klug and L.R. Silberbauer have been supported by the MDPhD Excellence Program of the Medical University of Vienna. M.B. Reed and L. Silberbauer were recipients of a DOC fellowship of the Austrian Academy of Sciences at the Department of Psychiatry and Psychotherapy, Medical University of Vienna. S. Jamadar is supported by a National Health and Medical Research Council of Australia Fellowship APP1174164. We thank the graduated team members and the diploma students of the Neuroimaging Lab (NIL, headed by R. Lanzenberger) as well as the clinical colleagues from the Department of Psychiatry and Psychotherapy of the Medical University of Vienna for clinical and/or administrative support. In detail, we would like to thank S. Kasper, K. Papageorgiou, P. Michenthaler, T. Vanicek, A. Basaran, M. Hienert, J. Unterholzner, G. Gryglewski and L. Rischka for medical or technical support, V. Ritter, K. Einenkel and E. Sittenberger for participant recruitment and A. Jelicic for partly implementation of the task. We are further grateful to J. Raitanen, J. Völkle, A. Pomberger, V. Pichler, W. Wadsak and the radioligand synthesis team from the Department of Biomedical Imaging and Image-guided Therapy, Division of Nuclear Medicine for acquisition support and supervision. The scientific project was performed with the support of the Medical Imaging Cluster of the Medical University of Vienna.

## Authors’ contributions (CRediT)

**Murray Bruce Reed**

Conceptualization, Formal Analysis, Data Curation, Methodology, Software, Visualization, Writing – Original Draft Preparation

**Magdalena Ponce de León**

Methodology, Writing – Review & Editing.

**Sebastian Klug**

Data Curation, Investigation, Writing – Review & Editing.

**Christian Milz**

Writing – Review & Editing.

**Leo Silberbauer**

Writing – Review & Editing.

**Pia Falb**

Writing – Review & Editing.

**Godber Mathis Godbersen**

Data Curation, Investigation, Writing – Review & Editing.

**Sharna Jamadar**

Software, Writing – Review & Editing.

**Zhaolin Chen**

Software, Writing – Review & Editing.

**Lukas Nics**

Project Administration, Writing – Review & Editing.

**Marcus Hacker**

Project Administration, Resources, Writing – Review & Editing.

**Rupert Lanzenberger**

Conceptualization, Resources, Supervision, Writing – Review & Editing.

**Andreas Hahn**

Conceptualization, Data Curation, Funding Acquisition, Investigation, Methodology, Project Administration, Supervision, Writing – Review & Editing.

All authors discussed the implications of the findings and approved the final version of the manuscript.

## Data Availability Statement

Raw data will not be publicly available due to reasons of data protection. Processed data and custom code can be obtained from the corresponding author with a data-sharing agreement, approved by the departments of legal affairs and data clearing of the Medical University of Vienna.

## Supplemental material

Supplemental material for this article is available online.

