## Supplementary Material for "Optimal filtering strategies for task-specific functional PET imaging"

***For submission to:***

***bioRxiv***

**Running Title: Optimal filters for fPET**

### Correspondence to:
Assoc. Prof. PD. Dr. Andreas Hahn, MSc

ORCID: https://orcid.org/0000-0001-9727-7580
Medical University of Vienna, Department of Psychiatry and Psychotherapy, Austria

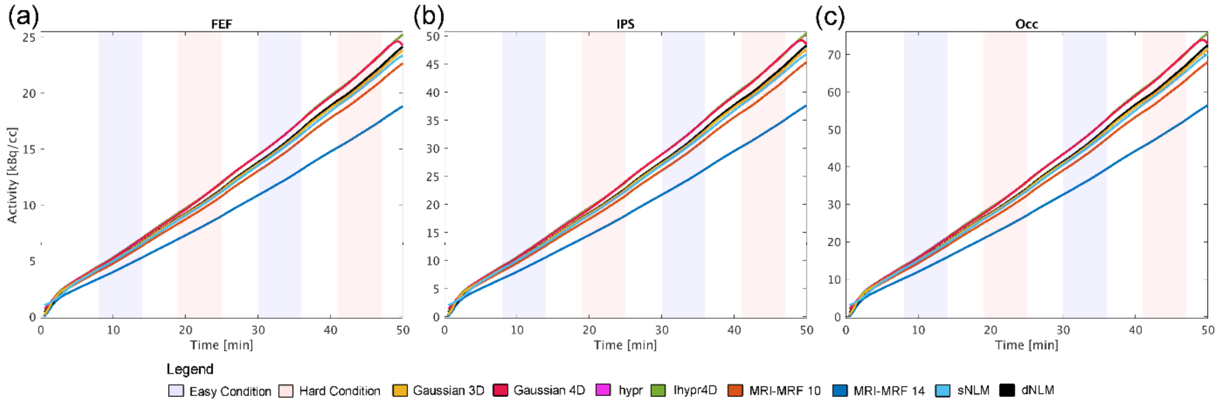


Supplementary Figure 1: Overview of time activity curves (averaged across participants) obtained with the different filtering techniques for a subset of participants whose task difficulty was ordered as easy-hard-easy-hard (n = 10), extracted from the (a) frontal eye field (FEF), (b) intraparietal sulcus (IPS) and (c) occipital cortex (OCC).

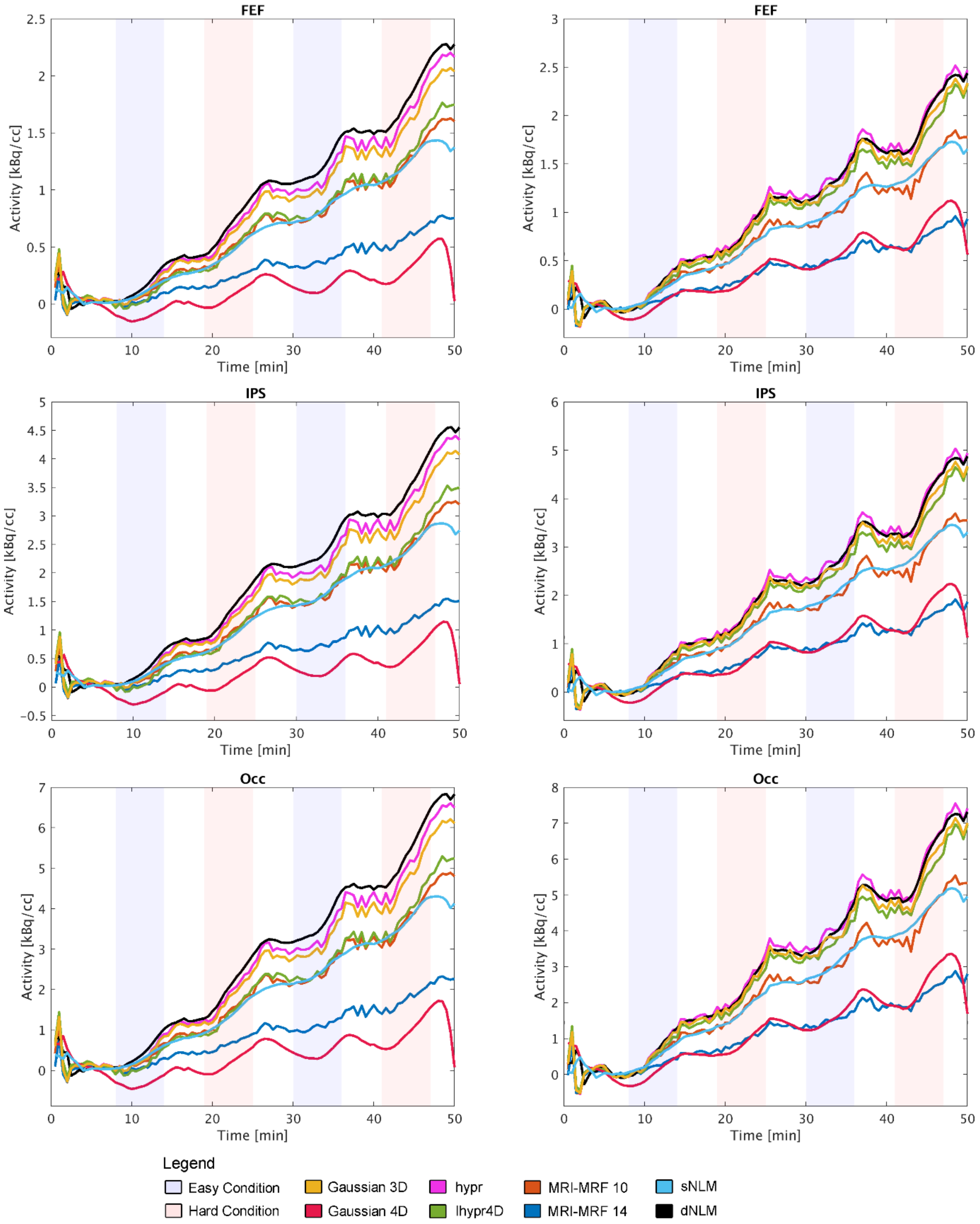


Supplementary Figure 2: Overview of task-specific regressors (Hard > BL) obtained with the different filtering techniques for two representative participants (left and right columns, respectively). Region of interest included the frontal eye field (FEF), intraparietal sulcus (IPS) and the occipital cortex (OCC).

| Filter | FEF | | IPS | | OCC | | Mean |
|---|---|---|---|---|---|---|---|
| | Easy | Hard | Easy | Hard | Easy | Hard | |
| **3D Gaussian** | 0.50 | 0.65 | 0.52 | 0.76 | 0.33 | 0.65 | 0.57 |
| **4D Gaussian** | 0.24 | 0.29 | 0.49 | 0.15 | 0.50 | 0.16 | 0.31 |
| **Hypr** | 0.51 | 0.66 | 0.53 | 0.77 | 0.33 | 0.65 | 0.58 |
| **Ihypr4D** | 0.40 | 0.50 | 0.46 | 0.70 | 0.22 | 0.62 | 0.48 |
| **MRI-MRF 10** | 0.47 | 0.64 | 0.52 | 0.75 | 0.30 | 0.60 | 0.55 |
| **MRI-MRF 14** | 0.44 | 0.66 | 0.48 | 0.70 | 0.09 | 0.46 | 0.47 |
| **Static NLM** | 0.43 | 0.56 | 0.34 | 0.78 | 0.27 | 0.63 | 0.50 |
| **Dynamic NLM** | 0.54 | 0.47 | 0.68 | 0.80 | 0.47 | 0.69 | 0.61 |

Supplementary Table 1: Detailed overview of Intraclass Correlation Coefficients separately for each region of interest and both task difficulties. The total average value is the same as in table 1 of the main text. These regions comprise the frontal eye field (FEF), intraparietal sulcus (IPS) and the occipital cortex (OCC).

| Filter | FEF | | IPS | | OCC | | Mean |
|---|---|---|---|---|---|---|---|
| | Easy | Hard | Easy | Hard | Easy | Hard | |
| **3D Gaussian** | 9.06 | 13.50 | 10.26 | 12.01 | 8.67 | 13.71 | 11.20 |
| **4D Gaussian** | 1.02 | 5.52 | 1.12 | 5.15 | 0.88 | 9.06 | 3.79 |
| **Hypr** | 8.95 | 13.32 | 10.25 | 12.04 | 8.64 | 13.74 | 11.16 |
| **Ihypr4D** | 9.18 | 15.61 | 10.62 | 12.69 | 8.28 | 13.93 | 11.72 |
| **MRI-MRF 10** | 8.81 | 13.72 | 10.39 | 12.29 | 8.87 | 13.74 | 11.30 |
| **MRI-MRF 14** | 7.54 | 12.35 | 9.40 | 11.79 | 8.74 | 13.38 | 10.53 |
| **Static NLM** | 12.68 | 15.14 | 13.1 | 14.28 | 11.19 | 14.52 | 13.49 |
| **Dynamic NLM** | 10.30 | 15.84 | 10.51 | 11.90 | 9.44 | 14.20 | 11.98 |

Supplementary Table 2: Detailed overview of temporal signal to noise ratio (tSNR) estimated separately for each region of interest and task difficulty. The total average value is the same as in table 1 of the main text. The three regions include the frontal eye field (FEF), intraparietal sulcus (IPS) and the occipital cortex (OCC).

| Filter | FEF | | IPS | | OCC | | Mean |
|---|---|---|---|---|---|---|---|
| | Easy | Hard | Easy | Hard | Easy | Hard | |
| **3D Gaussian** | 4.60 | 5.53 | 4.18 | 4.70 | 3.43 | 4.60 | 4.51 |
| **4D Gaussian** | 0.43 | 2.96 | 0.37 | 2.14 | -0.40 | 2.56 | 1.34 |
| **Hypr** | 4.51 | 5.43 | 4.14 | 4.64 | 3.38 | 4.53 | 4.44 |
| **Ihypr4D** | 4.59 | 5.65 | 4.18 | 4.79 | 3.39 | 4.69 | 4.37 |
| **MRI-MRF 10** | 4.54 | 5.61 | 4.48 | 5.00 | 3.92 | 5.26 | 4.80 |
| **MRI-MRF 14** | 5.06 | 6.87 | 5.58 | 6.32 | 5.43 | 7.44 | 6.12 |
| **Static NLM** | 8.12 | 10.30 | 7.48 | 8.40 | 6.86 | 9.34 | 7.58 |
| **Dynamic NLM** | 6.27 | 7.18 | 5.65 | 6.51 | 5.07 | 6.63 | 6.22 |

Supplementary Table 3: Detailed overview of the mean T-value extracted for each region of interest and task difficulty separately. The average value is the same as in table 1 of the main text. The three regions include the frontal eye field (FEF), intraparietal sulcus (IPS) and the occipital cortex (OCC).

| Filter | FEF | | IPS | | OCC | | Mean |
|---|---|---|---|---|---|---|---|
| | Easy | Hard | Easy | Hard | Easy | Hard | |
| **3D Gaussian** | 11.63 | 11.58 | 9.78 | 11.50 | 9.04 | 10.99 | 10.75 |
| **4D Gaussian** | 6.32 | 9.04 | 4.85 | 7.08 | 2.00 | 8.54 | 6.31 |
| **Hypr** | 10.76 | 11.66 | 9.43 | 11.03 | 8.65 | 11.15 | 10.45 |
| **Ihypr4D** | 11.88 | 11.70 | 11.86 | 12.02 | 7.78 | 12.16 | 11.24 |
| **MRI-MRF 10** | 12.30 | 13.65 | 11.05 | 12.25 | 8.93 | 12.97 | 11.86 |
| **MRI-MRF 14** | 9.97 | 15.26 | 11.96 | 12.54 | 10.75 | 18.23 | 13.12 |
| **Static NLM** | 12.84 | 16.59 | 13.69 | 16.31 | 11.97 | 21.18 | 15.57 |
| **Dynamic NLM** | 13.13 | 13.97 | 12.04 | 17.26 | 10.47 | 18.19 | 14.18 |

Supplementary Table 4: Detailed overview of the peak T-value extracted for each region of interest and task difficulty separately. The average value is the same as in table 1 of the main text. The three regions encompass the frontal eye field (FEF), intraparietal sulcus (IPS) and the occipital cortex (OCC).
